# Disentangling semantic and response learning effects in color-word contingency learning

**DOI:** 10.1101/546689

**Authors:** Sebastian Geukes, Dirk Vorberg, Pienie Zwitserlood

## Abstract

It is easier to indicate the ink color of a color-neutral noun when it is presented in the color in which it had been shown frequently before, relative to print colors in which it had been shown less often. This phenomenon is known as color-word-contingency learning. It remains unclear whether participants actually learn semantic (word-color) associations or/and response (word-button) associations. We here present a novel variant of the paradigm that can disentangle semantic and response learning, because word-color and word-button associations are manipulated independently. In four experiments, each involving four daily sessions, novel words (pseudowords such as *enas*, *fatu* or *imot*) were probabilistically associated either with a particular color, a particular response-button position, or with both. Neutral trials were also included, and participants’ awareness of the contingencies was manipulated. The data showed no impact of explicit contingency awareness, but clear evidence both for response learning and for semantic learning, with effects emerging swiftly. Deeper processing of color information, with color words presented in black instead of color patches to indicate response-button positions, resulted in stronger effects, both for semantic and response learning. Our data add a crucial piece of evidence lacking so far in color-word contingency learning studies: Semantic learning effectively takes place even when associations are learned in an incidental way.

## Introduction

Children learning their first language, and adults learning a foreign language, are permanently trying to establish reliable connections between words and the meaning they refer to. Learning word-concept relations is thus one of the core challenges associated with mastering a novel language, and by now there is ample evidence that such learning can be fast and efficient (e.g., Borovsky, Elman, & Kutas, 2012; Carey & Bartlett, 1978; Mestres-Missé, Rodríguez-Fornells, & Münte, 2007; McMurray, Kapnoula & Gaskell, 2016, for an overview).

To mimic that, in real life, information about potential word-concept relations is often ambiguous, many studies have employed a statistical learning approach: Novel words and their concepts are paired in a probabilistic manner, and learners must infer the correct word-concept relations over the course of the learning phase (Breitenstein et al., 2007; Breitenstein & Knecht, 2002; Dobel et al., 2010; Geukes, Gaskell, & Zwitserlood, 2015; Geukes & Zwitserlood, 2016; Smith & Yu, 2008; Yu & Smith, 2007). For example, Breitenstein et al. (2007) presented their participants with stimulus pairs consisting of an object picture and a spoken novel word (which were pseudowords in the participants’ native language). A particular word (e.g., *binu*) was paired with a particular object picture (e.g., a dog) half of the time (the to-be-learned combination of novel word and meaning), in the other half the word was combined with many different objects. As expected, telling apart correct from incorrect pairs – the task during learning - took time, but participants successfully learned to associate the semantics of concepts with correct novel words, as they could easily tell apart correct from incorrect pairs. Crucially, novel words (e.g., *binu*) effectively primed semantically related stimuli (e.g., picture of a cat), showing that they were indeed tightly connected to the conceptual semantic network.

Color concepts provide an interesting semantic field to study the association of novel words to existing meaning. Given that the conceptual space is small, it provides an ideal testing ground. Surprisingly, it has hardly been used (but see Altarriba & Mathis, 1997). In a previous study, we employed the statistical word-learning paradigm to test the association of novel words to color concepts. Effects of learning were assessed in terms of congruency effects in the manual Stroop task, in which the colors are identified by means of a key press (Geukes, Gaskell & Zwitserlood, 2015). Novel words that had been associated with native-language color words led to sizeable Stroop congruency effects on color-matching speed. This novel-word Stroop effect arose under two conditions: (1) immediately after learning, but only when novel words and native-language color words were present in the same Stroop blocks, or (2) with and without native-language color words, when tested 24 hours after learning, providing an opportunity for memory consolidation^i^.

Crucially, we had to make sure in this experiment that no further learning of the correct pair took place within the Stroop task. In fact, there is reliable evidence that the congruency effect in the manual Stroop task is partially due to word-response associations learned during the experiment itself (De Houwer, 2003). This is because, in the typical four-color Stroop experiment with 50% congruent and 50% incongruent trials, each color word appears three times as often in its congruent print color as in either of the three incongruent print colors. This allows participants to learn direct word-to-response associations, which speeds up manual reactions on congruent trials. The relevance of this type of contingency learning is seen best when color-unrelated words (e.g., move, wide, rest) are used instead of color words. This has been done in a paradigm called color-word contingency learning (Schmidt, Crump, Cheesman, & Besner, 2007; see also Musen & Squire, 1993; Pritchatt, 1968). In the manual variant of the Stroop task, the print color of stimulus words is indicated by pushing the matching colored response button. There were two crucial differences to the classic version with native-language color words: (1) color-unrelated words (e.g., *month*) were used, rather than color words (e.g., *red*); (2) these words were presented more often in one print color (e.g., 75% in red) than in any of the other colors. These experiments showed that participants do indeed adapt to arbitrary word-color contingencies, as evidenced by faster responses to words presented in their frequent color than in any of the other colors (Schmidt et al., 2007; Schmidt, Augustinova, & De Houwer, 2018; Schmidt, De Houwer, & Besner, 2010). These adaptation effects arise after just a few learning trials, remain stable in size, and are unlearned rapidly (Schmidt et al., 2010).

While adaptation effects to presentation frequencies are clear-cut, their explanation is not. At least two possibilities come to mind: (i) According to the *word-concept* (or semantic) *learning* hypothesis, congruency effects arise because words (novel and color-neutral existing words alike) become associated with color concepts, which facilitates selection of the correctly colored response button. (ii) According to the *word-response* learning hypothesis, congruency effects arise because the word is probabilistically paired with one particular *response* (the one with the word’s most frequent color), such that participants become faster over time to select the correct manual response to a given word. Thus, it is valid to attribute congruency effects in color-word contingency learning to semantic associations only if such effects do indeed arise via word-concept learning. One way to distinguish semantic from response learning is to map the four colors typically used in color-word contingency learning onto two (neutrally colored) response buttons. In this setup, each button represents two colors (e.g.: “if the word’s print color is either blue or yellow, press the left button”). This was realized in Experiment 4 of Schmidt et al. (2007), in which the authors distinguished between three types of trials:

*- response mismatch*, when neither the print color nor the required response is the one most frequently associated with the presented word;

*- response match*, when the response but not the print color is associated with the word;

*- stimulus match*, when both print color and response are associated with the presented word.

These conditions allow distinguishing two separate learning components from each other: If there is *response learning*, responses should be faster on response match than on response mismatch trials. This is so because the word in the response match condition, while printed in a non-contingent color, nevertheless points to its correct response, thereby facilitating response selection. If there is *semantic learning*, however, responding should be faster on stimulus match than on response match trials. This is because the congruency between word and print color may facilitate the selection of the correct response over and above word-response congruency.

Schmidt et al. (2007) found evidence for response learning (a 26 ms effect), but no reliable advantage for stimulus-match over response-match trials (a net difference of 2 ms). Their conclusion was that color-word contingency learning reflects response learning only, not semantic learning. This is different in the classic manual Stroop task with native-language color words and a two-on-one response mapping, where both semantic and response congruency effects are reliably observed (Chen, Bailey, Tiernan, & West, 2011; De Houwer, 2003; Schmidt & Cheesman, 2005; van Veen & Carter, 2005).

While certainly informative, the 2:1 mapping task design has drawbacks. For example, while response options are reduced to two responses, there are still four colors to be distinguished, which could facilitate learning the response contingencies via color contingencies. Also, words are always correlated with both color and response. Consequently, semantic and response learning are neither learned nor tested independently, using separate sets of words. It is conceivable that participants do in fact learn word-color contingencies, which need not show up in response times, if the response contingency dominates any semantic (word-color) contingency effect. Semantic contingencies might lead to measurable effects, either (i) when learned independently of response-position contingencies, or (ii) when tested in different semantic paradigms. Thus, while the 2:1 mapping design does offer some evidence for dominant response-learning effects, it cannot be logically excluded that participants learn word semantics as well. A more general factor that might hamper semantic learning in the above studies is the fact that the words used possess meaning. Exactly because “move” or “rest” are not associated with a particular color – a prerequisite for their choice as stimuli – adding color to their semantic representation may be quite difficult to learn.

The goal of the current study was to separate semantic from response components in color-word contingency learning, by independently manipulating word-color and word-response associations across separate sets of novel words. To do so, we introduced a variant of the color-matching Stroop task in which the print color of novel words is to be reported via button presses, as in previous color-word contingency studies. We use pseudowords (e.g., *ugir*, *inwa*) which, by definition, are not associated with any color, as they have no established referential meaning at all. We refer to these stimuli as “pseudowords”, “novel words” or just “words”. The key difference from previous studies is the following: Simultaneously with the colored novel word, four color squares (Experiments 1, 2, and 4) are presented. The participants’ task is to indicate by button press which of the squares matches the word’s print color^ii^ (see Figure 1A). This allows us to vary the association of colors (red, green, blue or yellow) to response positions (button 1, 2, 3 or 4) on a trial-by-trial basis, thus permitting to separately manipulate word-color and word-position contingencies. In Experiment 1, 2 and 3, the assignment of a particular pseudoword to a particular contingency was held constant throughout; in Experiment 4 this assignment was re-randomized from block to block. (In Experiment 3, we replaced the colored squares by color words printed black, which surrounded the colored novel word; see Figure 6 below).

**Figure 1.**
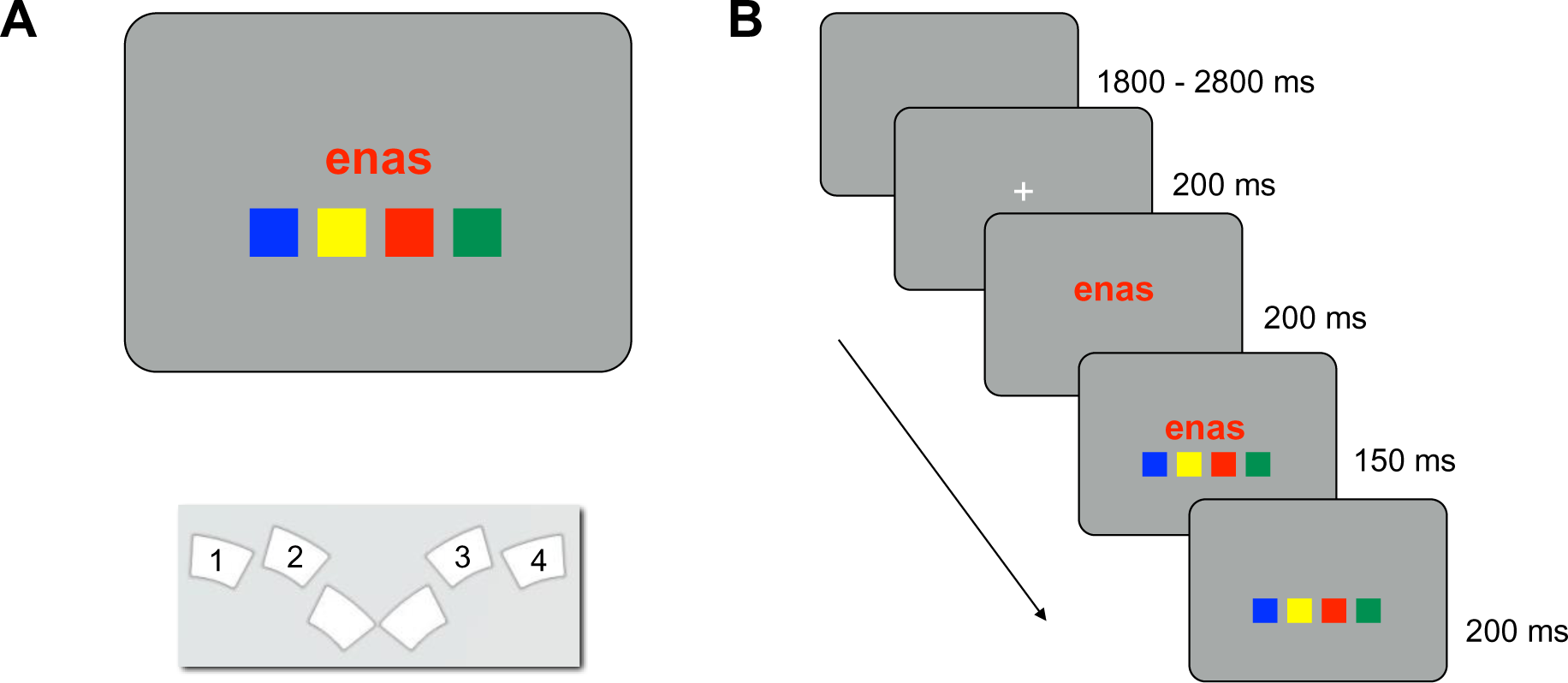
Stimulus display, spatial arrangement of response buttons, and trial timing. Panel A shows the typical stimulus display and response button layout in Experiments 1, 2, and 4. Panel B illustrates a typical trial. Numbers in Panel A schematically indicate which buttons of the response box were used in the experiments; actually, buttons were not labelled. The same timing of stimulus elements applied in all four experiments.

We used this paradigm in four experiments, to separate semantic from response learning effects in color-word contingency learning. Contingency type was blocked, with particular words associated either with color (Col), with spatial response position (Pos), or with both (ColPos). In a fourth contingency type, words were correlated with both color and response position, while the spatial order of the color patches was held constant throughout the experiment (ColPosFix), which closely resembles experiments with a spatially fixed set of colored response buttons.

Each experiment comprised four daily sessions, with two blocks of each contingency type in each session. The particular contingencies were determined randomly for individual participants but were kept constant throughout their experiment.

In Experiment 1, we tested whether the color-word contingencies do in fact induce congruency effects, and if so, how they depend on contingency conditions. In Experiment 2, we tested whether knowledge of the type of contingency present in the upcoming block changes the pattern of effects, as previous evidence suggests that explicit contingency awareness affects the size of congruency effects (Schmidt & De Houwer, 2012a). In Experiment 3, we used color words (printed in black) rather than color patches as response cues to indicate the current trial’s response mapping. Experiment 4 tested how rapidly contingency effects arise, by varying color-word contingencies from block to block, rather than keeping them constant across blocks.

Our predictions were as follows. We expected congruency effects in the ColPosFix condition that is closest to the study by Schmidt et al. (2007). We also predicted congruency effects in the condition that combines position and color information (ColPos). If color-word contingency learning reflects response learning, as Schmidt and colleagues claim, we predicted small but reliable congruency effects when only position information is present (Pos). Finally, we also expected evidence for semantic learning with color information in the absence of position information, because such information can become associated with pseudowords if cues are sufficiently consistent (Geukes et al., 2015). However, the effects should be smaller than with fixed spatial arrangement of the colored response cues, if ‘position habits’ are easily formed in learning, as suggested by findings in animal discrimination learning (e.g., Restle, 1962; Spence, 1956).

## Experimental 1

### Methods

#### Participants

Ten students (7 female; age: *M* = 23.50, *SD* = 3.14) took part, in four sessions on consecutive days. All had normal or corrected-to-normal visual acuity and normal color vision, as assessed by standard Ishihara plates. Participants gave written consent and received either course credit or 25 € compensation. All procedures were approved by the Ethics committee of the Department of Psychology, University of Münster.

#### Word materials

Twenty bi-syllabic pseudowords were used as stimuli (*ahak, alep, edok, emgu, enas, fatu, fepa, imot, inwa, kela, kopu, meha, mupa, palo, osig, ovon, sego, tihe, ugir, utaf*). They were selected from the materials developed and empirically validated by Breitenstein and Knecht (2002), who have shown that these pseudowords elicit few associations to existing German words and are of neutral emotional valence. On each trial, a single pseudoword, uniformly colored, was presented centered on a grey screen (see Figure 1), printed either red, green, blue, or yellow. RGB values: red (255, 0, 0), green (0, 255, 0), blue (255, 255, 0), yellow (0, 0, 255), grey (35, 35, 35).

#### Color matching task

A colored pseudoword (in red, green, blue, or yellow) appeared in the screen center, above a row of four differently colored squares, one of which always matched the word’s print color (see Figure 1A). The task was to locate the color-matching square, and to push the button corresponding to its position on the response box, as quickly as possible. Depending on experimental conditions, the spatial arrangement of the color squares on the screen either varied from trial to trial or was constant within a block (see below).

#### Contingency conditions

There were four contingency types. In 80% of the cases, a novel word either appeared (a) in one and the same print color (but at different positions; *Col*), (b) was associated with the same spatial response position (but varied in color; *Pos*), (c) had the same color and response position (*ColPos*) or (d) had the same color and response position, with a constant spatial order of the response cues within the block (*ColPosFix*).

For each participant, the 20 pseudowords were randomly partitioned into four subsets of five words each, and each subset was assigned to one of four contingency conditions. In each subset, one of the five words served as control, with print color and position of the matching color patch chosen randomly on any trial. The other four pseudowords were assigned to a contingency condition. For example, in a *Col*-contingency block, each of the four pseudowords was presented on 15 trials altogether, on 12 trials (p =.8) in its assigned color (‘congruent’, e.g. *blue*), and on 3 trials (1 - p = .2) in either one of the remaining three colors (‘incongruent’, e.g. *red, green*, or *yellow*). The same principle was used in *Pos*, Col*Pos* and *ColPosFix* blocks: *Pos*-words were assigned to one of the four response positions. They could appear in any color, but the correct response, that is, the spatial position of the matching color patch, was predictable (p=.8). Analogously, for *ColPos-words* and for *ColPosFix-words*, both color and response position were predictable (the latter with a fixed color-to-button assignment, see below).

**Figure 2.**
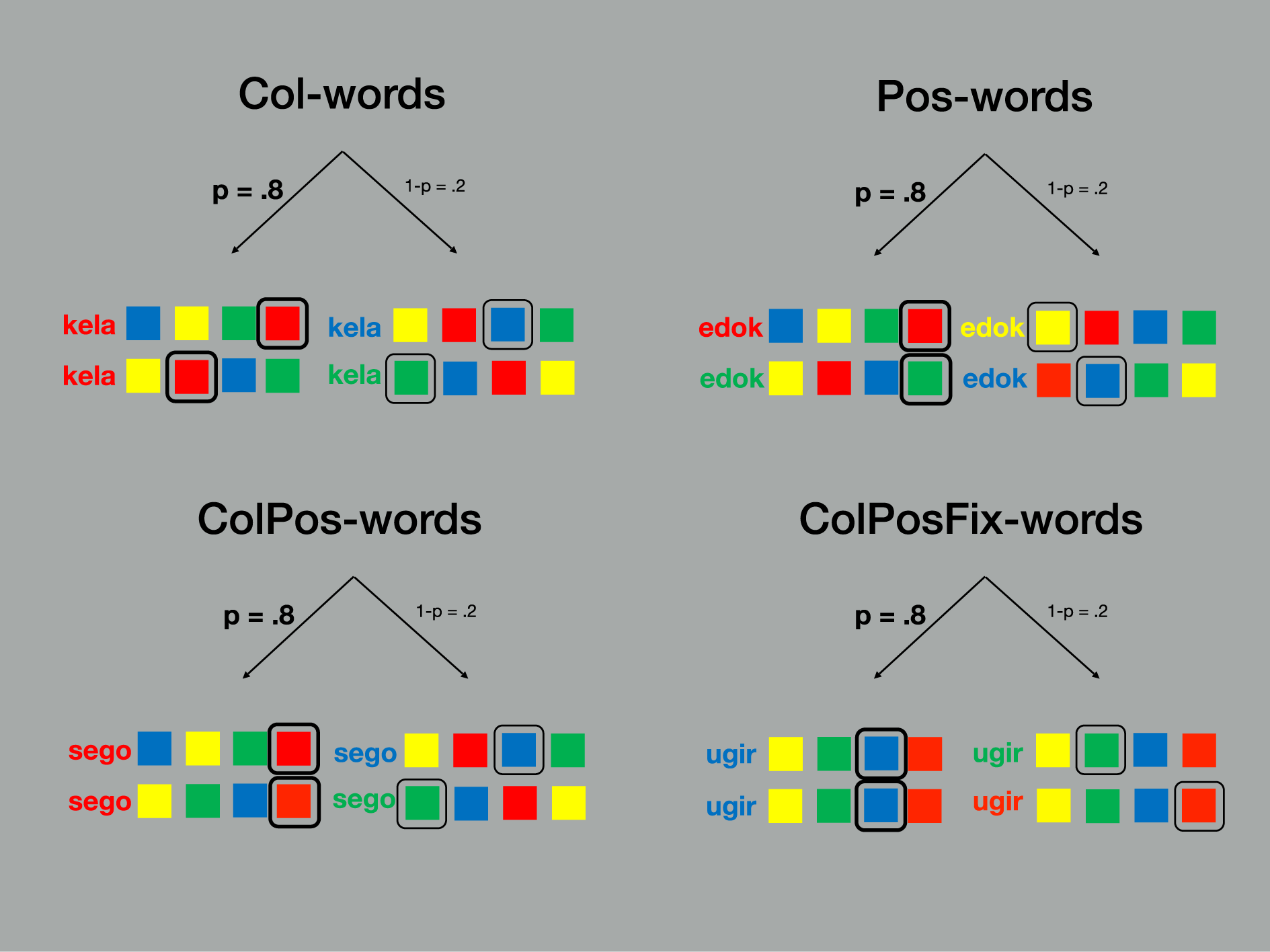
Four Contingency types. The figure illustrates how pseudowords were associated with color and/or response position in the Stroop task, yielding congruent (p = .8) and incongruent (1–p = .2) trials, where p denotes the probability of a congruent trial for a given word. For each contingency condition, two example trials are shown. For neutral words (not shown here), print color and response position were determined randomly on each trial. Note that for ColPosFix trials, the spatial order of the color squares remained fixed throughout, in contrast to ColPos trials, on which the non-target positions varied randomly.

#### Experimental conditions

The experiment followed a 4 x 3 x 8 experimental design: Contingency Type (*Col, Pos, ColPos, ColPosFix*) x Congruency (congruent, incongruent, neutral) x Block Repetition (1 to 8). Within each block of 75 trials, a fixed subset of five pseudowords was used: one neutral control word and four experimental words of the same type (either *Col, Pos, ColPos, or ColPosFix*). Each word appeared as colored stimulus on 15 trials per block. For experimental words, color and/or position of the matching color patch were consistent on 12 trials and inconsistent on 3 trials within each block. For neutral words, color and response position of the matching color patch varied randomly from trial to trial. As stated above, *ColPos*-words and *ColPosFix*-words were associated both with a color and a response position. *ColPos*-blocks and *ColPosFix*-blocks were constructed identically, except for the spatial arrangement of the color patches on the screen (see Figure 2). In *ColPos*-blocks, the three incorrect color patches appeared randomly at the remaining positions. in *ColPosFix*-blocks, the color patches were presented in a fixed spatial order throughout the entire block, analogously to a physically fixed arrangement of color-labelled response buttons.

#### Trial events and task

Stimuli were presented on an LCD color monitor (Samsung 2233 RZ, 120 Hz, 22 inch) using the *Presentation* software package (Neurobehavioral Systems, Inc.). Each trial started with a fixation mark, followed by a colored stimulus word and the four color patches. The exact timing of stimulus events is shown in Figure 1B.

Participants were to press the response button that spatially corresponded to the location of the color patch that matched the pseudoword’s color. (Response box: RB-830, Cedrus Corp., San Pedro, CA; see Figure 1A for the button layout). To make the particular contingencies less obvious and to avoid associations of particular visual stimuli to color and/or position, we randomly varied the font type (8 styles) and font size (4 sizes) of the stimulus words from trial to trial (see also Schmidt & Lemercier, 2018). Response times were measured from word onset. Incorrect responses or response omissions (response time > 2000 ms) were signaled by a low or high pitch tone, respectively, delivered via headphones.

Before the first main block, participants were given two practice blocks of 40 trials each. Each participant took part in four daily 1-h sessions, each of which contained eight contingency blocks with 75 trials. Blocks 1-4, and 5-8, contained each contingency type (*Col, Pos, ColPos, ColPosFix*); their order was random. Thus, each contingency type occurred twice per session, eight times overall. For each participant, individual assignments of pseudowords to within- and between-block conditions applied throughout the whole experiment. Performance feedback (mean response time, percentage correct) was provided after each block.

There was an obligatory 5-minutes rest after block 4. Participants were encouraged to also take rests between blocks if needed. Apart from the mid-session break, they started blocks at their discretion. No information was given about the task structure and its changes across trial blocks.

#### Analysis

For the response-time analyses, trials with RTs below 100 ms (less than 0.1% of all responses) were excluded, as well as error trials. For the remaining correct response trials, trimmed response-time means (5% trim at each distribution end, see Wilcox & Keselman, 2003) were calculated per participant and *Contingency type* (*Col, Pos, ColPos, ColPosFix*), *Congruency* (congruent, incongruent, neutral) and *Block Repetition* (1 to 8). Error rates were similarly calculated per experimental cell and subsequently arcsine square-root transformed. Trimmed response-time means and transformed error rates were analyzed by repeated-measures analysis of variance (ANOVA).

#### Awareness of contingencies

Following the final session, participants were handed a stack of cards each of which contained one of the 20 novel words printed in black. In two separate matching trials, participants were to match each word as best as possible both to (a) one of the four colors and (b) to one of the response positions. For this purpose, sheets were provided with four columns labelled either for the four colors or for the four response positions.

We determined the number of correct matches for the experimental words of each contingency type (*Col, Pos, ColPos, ColPosFix*), separately for color and for position contingency. For each contingency type, participants could correctly match between 0 and 4 words. Whether matching was better than chance (25%) was tested by one-sided one-sample *t*-tests. Considering the problems associated with confirming the null hypothesis in Null-Hypothesis Significance Testing, we also calculated JZS Bayes factors, assuming a standard Cauchy prior with r equal 0.707, using the JASP software package (Love et al., 2015).

### Results

Mean response times from Experiment 1 are shown in Figure 3A. Because the overall range of response times was large and there were considerable practice effects across sessions, the experimental effects of main interest are hard to discern from the absolute response times. Therefore, we include additional graphs of the net effects that show the differences between experimental conditions (Figure 3B).

Results from Experiment 1 showed a small but reliable difference of, on average, 12 ms between the congruent and incongruent conditions (this difference is henceforth called Congruency effect). This effect emerged early and remained relatively stable throughout the eight blocks. Response times to neutral words were in between those in congruent and incongruent conditions, suggesting the presence of both facilitation and inhibition. The ANOVA on response times revealed significant main effects of *Block Repetition*, *Contingency Type* and *Congruency*, but no reliable interactions. The *Congruency* main effect, *F*(2, 18) = 24.04, *p* < .001, was driven by faster responses on congruent (557 ms) than on neutral trials (566 ms), *t*(9) = 4.44, *p* = .002, which in turn were faster than those on incongruent trials (569 ms), *t*(9) = 3.04, *p* = .013. Response times also differed between *Contingency Types*, *F*(3, 27) = 46.09, *p* < .001; responses were slowest in *Pos*-blocks (593 ms), followed by *Col*-blocks (586 ms), *ColPos*-blocks (547 ms) and *ColPosFix*-blocks (530 ms). All differences between Contingency Type mean response times were significant with *t*(9) ≥ 3.69, *p* ≤ .005. Finally, response speed improved with *Block Repetition*, *F*(7, 63) *=* 10.92, *p* < .001, decreasing from 591 ms to 549 ms over the eight blocks. Of the interactions, the *Congruency* × *Contingency Type* interaction just failed significance, *F*(6, 54) = 2.09, *p* = .069.

To test our predictions for the four contingency conditions, follow-up tests showed that the response-time difference between congruent and incongruent trials failed significance in the *Col*-condition (4 ms, *t*(9) = 1.33, *p* = 0.216), but was significant in the three conditions with position contingencies (*Pos*: 14 ms, *t*(9) = 2.80, *p* = 0.021; *ColPos*: 15 ms, *t*(9) = 4.35, *p* = 0.002; *ColPosFix*: 15 ms, *t*(9) = 8.57, *p* < .001). None of the remaining interactions turned out reliable (*Block* × *Contingency type*: *F*(21, 189) = 1.32, *p* = .160; *Congruency* × *Block Repetition*: *F*(14, 126) = 0.93, *p* = .532; *Congruency* × *Block Repetition* × *Contingency Type*, *F*(42, 378) = 0.91, *p* = .638).

**Figure 3.**
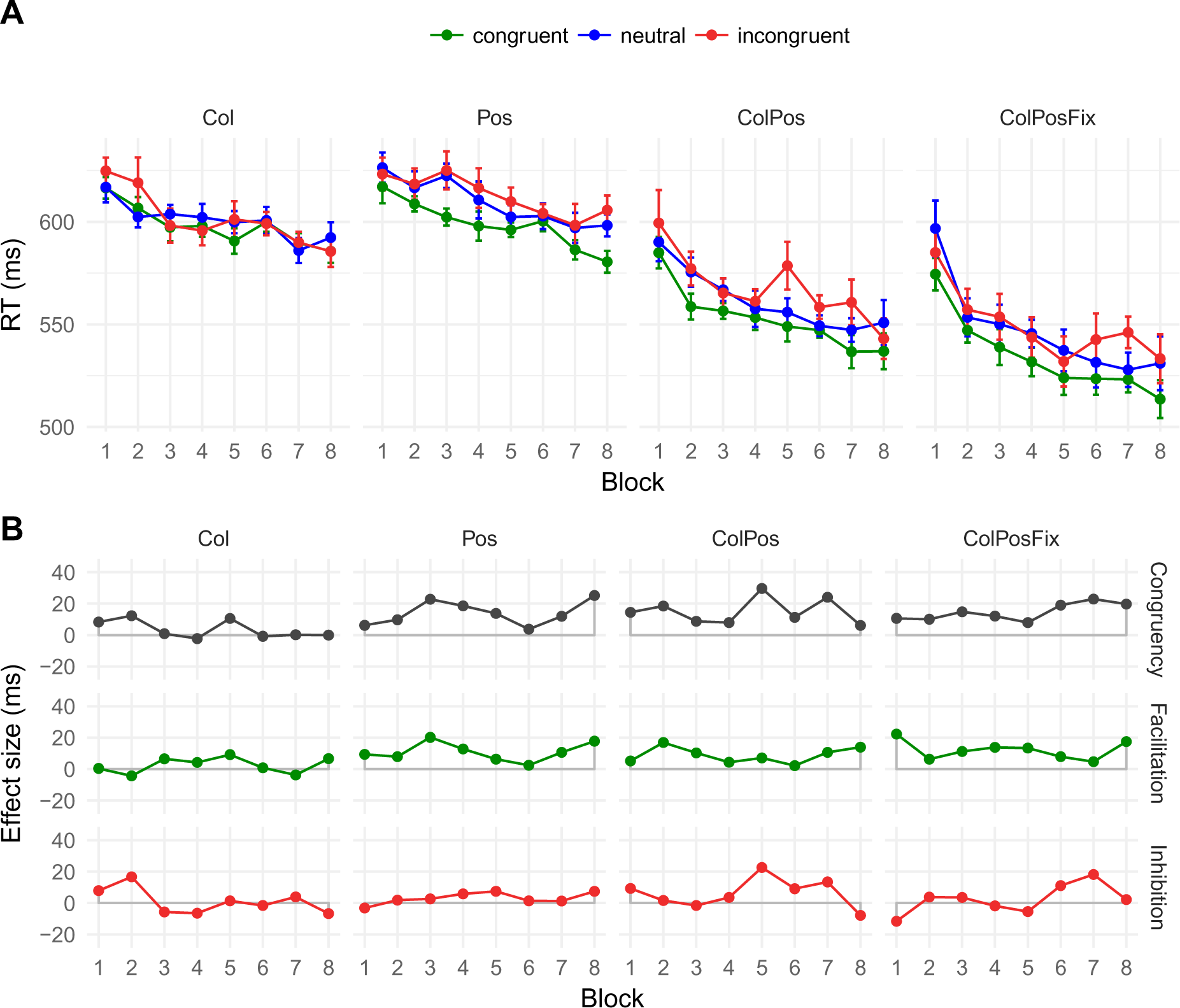
Mean response times (A) and net effects (B) in Experiment 1 (n = 10). Error bars are within-participant standard errors of the mean 29,30. Abbreviations of the Contingency Types: Col = color contingency, Pos = response position contingency, ColPos = both contingencies, ColPosFix = both contingencies plus fixed spatial arrangement of the response cues (see Methods for details). Effects: Congruency = RT_incongruent_ – RT_congruent_, Facilitation = RT_neutral_ – RT_congruent_, Inhibition = RT_incongruent_ – RT_neutral_

Error rates were low, around 3%, and the pattern of error rates across congruency conditions was roughly consistent with the pattern in response times (congruent: 2.7%, neutral: 3.1%, incongruent: 3.1%; see Supplementary Materials, Figure A1). The ANOVA of the (arcsine square-root transformed) error rates yielded a marginal main effect of *Congruency, F*(2, 18) = 3.50, *p* = .052. This congruency pattern was present in all blocks with position contingencies (*Pos*: congruent 2.7%, incongruent 4.2%; *ColPos*: congruent 2.4%, incongruent 3.1%; *ColPosFix*: congruent 2.2%, incongruent 2.8%), but it was reversed in the *Col*-condition (congruent 3.3%, incongruent 2.3%). There was also a marginal three-way interaction of *Congruency*, *Block Repetition*, and *Contingency Type*, *F*(42, 378) = 1.42, *p* = .050. No other interactions or main effects were statistically reliable (all *F*s < 1.86, all *p* > .105).

#### Contingency Awareness

Unfortunately, the files containing the descriptive statistics for the matching task were lost, but the originally calculated inferential statistics are available in the diploma thesis that also reported this experiment (Praast, 2013). These offer the necessary information about whether participants could match words to colors and words to positions better than chance. With words from the *ColPosFix* contingency, participants could match words both to colors (*t*[9] = 2.86, *p* = .009, BF_10_ = 3.76,iii) and to response positions (*t*[9] = 3.10, *p* = .007, BF_10_ = 5.11) with above chance performance, but the Bayes Factors indicate only moderate evidence for this. For the *Pos* contingency, matching words to positions was not better than chance (*t*[9] = 0.43, *p* = .338, BF_01_ = 2.99). In the other conditions, Bayes Factors indicated that the data are inconclusive (BF_10_ close to 1; see Supplementary Materials, Table A1, for details).

### Discussion

Experiment 1 employed a novel paradigm to assess the emergence of semantic- and response-congruency effects during color-word-contingency learning. Overall, participants performed with sufficient ease and speed in this rather complex task. Responses were faster when both color and position were correlated with the stimulus words (*ColPos*-blocks), compared to conditions under which one dimension only was validly coupled with the words (*Col*- and *Pos*-blocks). As expected, responses were even faster when response positions remained constant across the entire block (*ColPosFix*- blocks). This pattern suggests that random variation in the presentation – either in the uncorrelated dimensions (*Col*- and *Pos*-blocks) or in the positioning of the non-relevant response cues (*ColPos*-blocks) – slows responding, as compared to blocks in which both dimensions are correlated and the color-to-response mapping is constant (*ColPosFix*- blocks).

Crucially, there was a statistically reliable congruency effect in response times, suggesting that participants were able to adapt behavior to the implemented stimulus-response contingencies. Blocks with position contingencies (*Pos*, *ColPos* and *ColPosFix*) yielded clear congruency effects, but the effects were numerically small and missed significance in color contingency blocks (see Figure 3). This is in line with previous color-word-contingency learning studies (Schmidt et al., 2007), but, considering the relatively small congruency effects, the current design seems not sufficiently powerful to corroborate the apparent interaction of congruency and contingency type. Interestingly, congruency effects emerged very early but did not substantially change in size over the course of no less than eight blocks, which is again consistent with previous findings (Schmidt et al., 2010). It thus seems that the underlying learning processes are fast, but, at least in our complex experimental setup, adaptation to the particular contingencies is limited to relatively small gains of about 10 to 20 ms.

In one of their studies on color-word-contingency learning, Schmidt and De Houwer (2012a) tested whether the size of the congruency effect depends on participants having explicit information about the *general underlying contingency* (e.g., “each word in the experiment is presented most often in a certain color”), paired with the instruction to learn the contingency. Using three color-word pairs, congruency effects were slightly larger when participants had been informed about the word-color contingency than when not. Schmidt and De Houwer (2012b) also tested whether revealing the *specific word-color pairs* (e.g., “the word ‘month’ is presented most often in red”) had an impact on the congruency effect and reported small but just significant effects in favor of this hypothesis. The aim of Experiment 2 was to test whether awareness of the contingencies affects the response-time pattern in our more complex setup.

## Experiment 2

To check the impact of explicit knowledge of the contingencies, we re-ran Experiment 1, this time providing information about the *general type of contingency* (Color, Position, Color & Position) implemented in each upcoming block (similar to De Houwer, 2012a). As in Experiment 1, we assessed awareness of the specific contingencies by having participants match the stimulus words either to the four colors, or the four response positions.

### Methods

#### Participants

Twelve students (5 male, age: *M* = 26.17, *SD* = 4.95) took part who had not participated in Experiment 1. Inclusion criteria, informed consent, and compensation were the same as in Experiment 1.

#### Procedure

Experiment 2 was identical to Experiment 1, except that, before each block, participants were informed about the contingencies to be expected in the following trials. For example, before color-contingency blocks, participants read the statement: “BLOCKTYPE 1: COLORS. In the following block, you will see five different words. Four of these will frequently appear in a particular COLOR. Thus, the word can guide you to find the correct response color” (translated from the German instruction). Equivalent information was presented for each contingency type. Thus, participants were given the opportunity to observe and profit from the particular contingencies. In the final matching task, the neutral (uncorrelated) words of the different contingency types were now included and a “neutral” matching option was provided on the response sheets. The chance level was therefore at 20%.

#### Analysis

Response times and error rates were analyzed as above.

### Results

Mean response times and net congruency effects are presented in Figure 4. The pattern was very similar to that of Experiment 1: There was an 11 ms difference between congruent and incongruent trials (Exp. 1: 12 ms). The ANOVA on response times revealed a significant *Congruency* main effect, *F*(2, 22) = 22.42, *p* < .001, with overall faster responses on congruent (583 ms) than on neutral trials (591 ms), which in turn were faster on incongruent trials (594 ms). The mean difference between congruent and to neutral trials was small but reliable (8 ms, *t*(11) = 6.08, *p* < .001); this was not the case for the difference between neutral and incongruent (3 ms), *t*(11) = 1.39, *p* = .193. Mean response times again decreased with *Block Repetitions*, *F*(7, 77) *=* 21.25, *p* < .001, from 624 ms in Block 1 to 570 ms in Block 8. Finally, response times depended on Contingency *Type*, *F*(3, 33) = 87.64, *p* < .001, with the same pattern as in Experiment 1: Responses were slowest in *Pos*-blocks and in *Col*-blocks (both means 618 ms), faster in *ColPos* blocks (576 ms) and *ColPosFix* blocks (546 ms). Response time means in *Col*- and *Pos*-blocks did not reliably differ, *t*(11) = 0.27, *p* = .800, but mean *Col*-RT differed from *ColPos*-RT by 42 ms, *t*(11) = 11.05, *p* < .001, and mean *ColPos*-RT differed from *ColPosFix* RT by 30 ms, *t*(11) = 5.31, *p* < .001. Although none of the interactions was close to significance (all *F* < 1.50, *p* > .192), indicating congruency in all contingency conditions, we again tested our predictions for the different contingency types by separately assessing c*ongruency* effects. For all Contingency Types, responding on congruent trials was significantly faster than on incongruent trials (*Col*: 7 ms, *t*(11) = 2.25, *p* = .046, *Pos*: 8 ms, *t*(11) = 2.66, *p* = .022, *ColPos*: 17 ms, *t*(11) = 4.42, *p* = .001, *ColPosFix*: 11 ms, *t*(11) = 3.07, *p* = .011).

**Figure 4.**
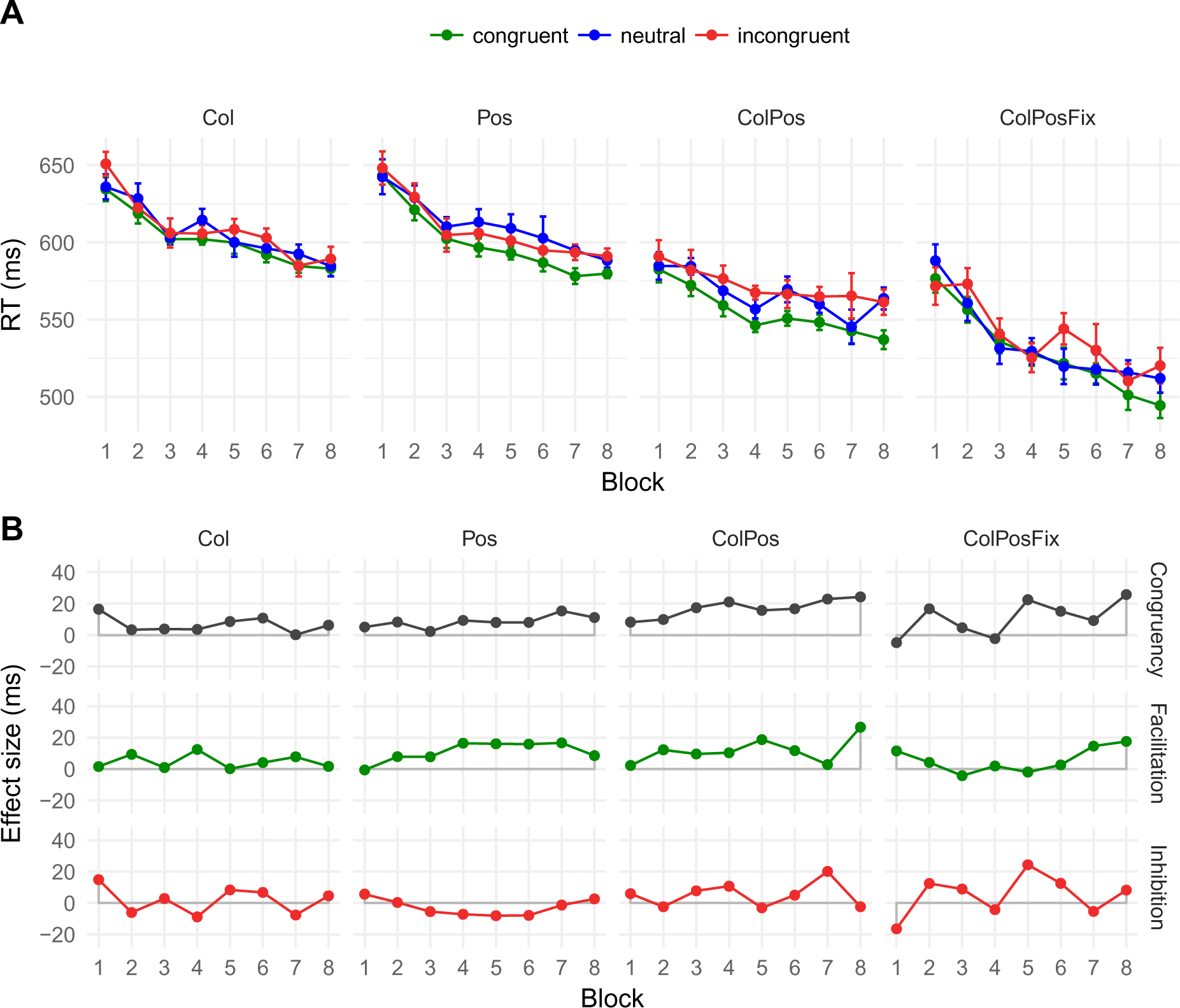
Mean response times (A) and net effects (B) in Experiment 2 (n = 12). In contrast to Experiment 1, participants were made aware of the contingency type before each upcoming block. Error bars and abbreviations as in Figure 3.

The error rate amounted to 3% of all trials and showed the pattern expected from congruency (congruent 2.6%, neutral 2.9%, incongruent 3.7%; see Supplementary Materials, Figure A1); however, this *Congruency* effect failed significance, *F*(2, 22) = 1.82, *p* = .186, as did the other main and interaction effects (*Contingency Type* main effect: *F*(2, 22) = 2.76, *p* = .057, all other *F* ≤ 1.12, *p* ≥ .361).

#### Contingency Awareness

In the matching task that followed the last block, participants performed better than chance when matching words from the *ColPosFix* block to colors, *M* = 42%, *SD* = 26%, *t*(11) = 2.86, *p* = .008, BF_10_ = 8.23. Despite congruency instructions, performance levels were close to chance under all remaining conditions (range of % correct matches: 18 – 28% [chance level = 20%], all *t*[11] < 1.24, *p* > .121) and either inconclusive or in favor of the null (BF_01_ between 1.08 and 3.48; see Supplementary Materials, Table A2, for details).

#### Joint Analysis of Experiments 1 and 2

Because the different instructions in Experiments 1 and 2 did not seem to affect the pattern of latency effects, we ran a joint post-hoc analysis of the two experiments. The ANOVA of the combined data with *Experiment* as between-participants factor revealed no reliable effects of this factor (main effect: *F*(1, 20) = 2.43, *p* = .135; all interactions with *Experiment*: *F* < 1.33, all *p* > .273). We therefore ran a joint ANOVA, collapsing the data from both experiments (Figure 5) to increase power. Again, all three main effects turned out clearly significant, confirming the pattern from the individual experiments (all *F* > 31.33, all *p* < .001). As before, most of the interactions turned out non-significant (*Block Repetition* × *Contingency Type: F*[21, 441] = 1.34, *p* = .145; *Congruency × Block Repetition*, *F*[14, 294] = 0.92, *p* = .536; *Congruency × Block Repetition × Contingency Type: F*[42, 882] = 0.95, *p* = .562). However, there was now a significant *Congruency* by *Contingency Type* interaction, *F*(6, 126) = 2.61, *p* = .020, suggesting that the size of the congruency effect was affected by the type of contingency. Indeed, although congruency effects were reliable for all contingency types, they clearly differed in size: *Col*: 5 ms (*t*[21] = 2.62, *p* = .016); *Pos*: 8 ms (*t*[21] = 3.03, *p* = .006), *ColPos*: 13 ms (*t*[21] = 5.52, *p* < .001) and *ColPosFix*: 11 ms (*t*[21] = 5.29, *p* < .001).

**Figure 5.**
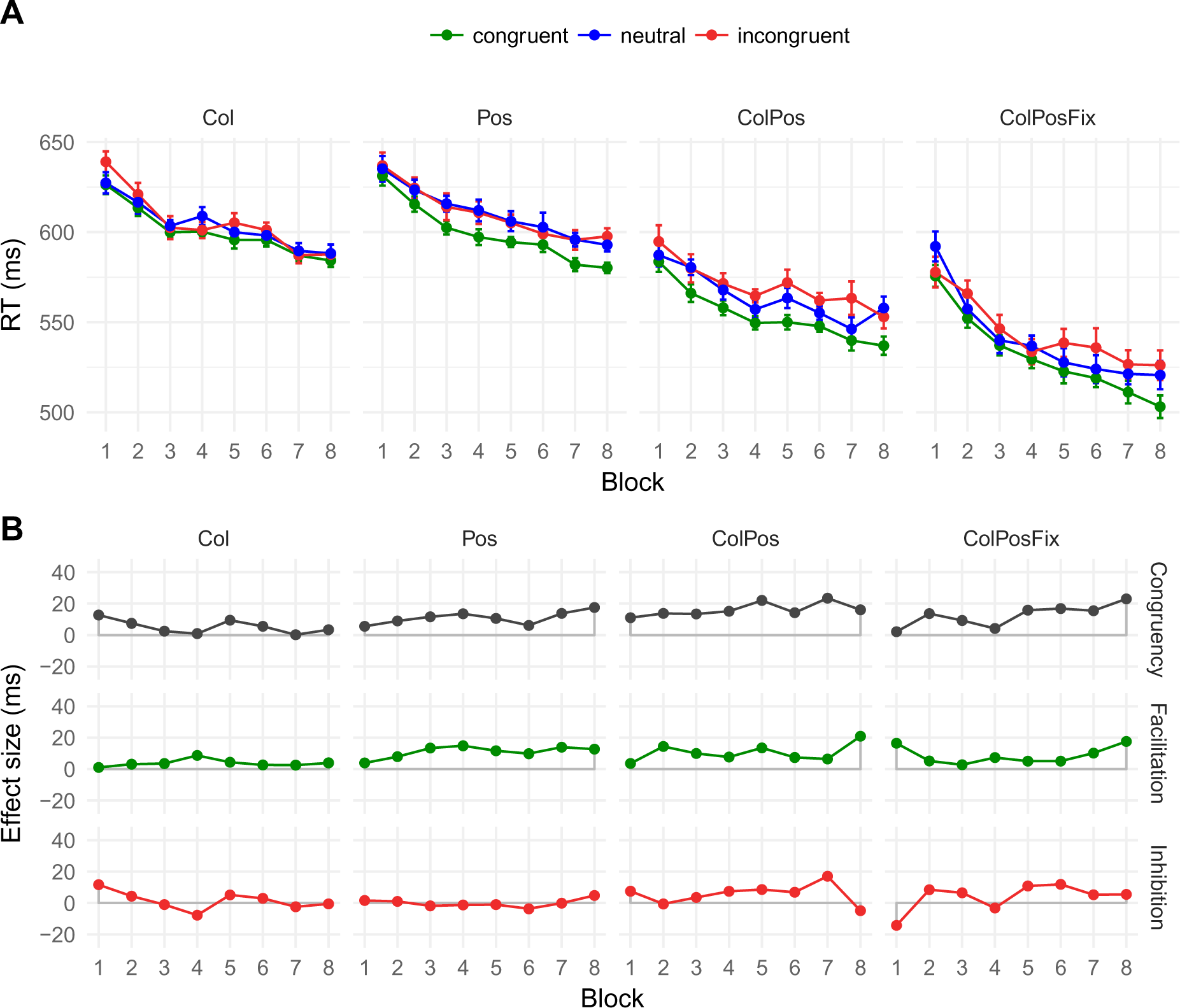
Mean response times (A) and net effects (B), collapsed over Experiments 1 and 2 (combined n = 22). Error bars and abbreviations as in Figure 3.

The error rates from Experiments 1 and 2 combined followed the pattern expected from *Congruency*, with congruent, neutral, and incongruent trials leading to 2.61%, 3.00%, and 3.41% of errors, respectively. This was confirmed by a significant main effect of *Congruency*, *F*(2, 42) = 4.11, *p* = .024. None of the other main or interaction effects reached significance (all *F* ≤ 1.62, all *p* ≥ .133).

### Discussion

Experiment 2 differed from Experiment 1 only in that participants were informed before each block about the contingencies in the upcoming trials. Despite this explicit instruction, participants were afterwards unable to match words to colors and/or to positions, except for contingencies of the *ColPosFix* blocks, which also showed above-chance matching in Experiment 1. Latency effects were virtually identical in Experiments 1 and 2, and their joint analysis provided no evidence that the different instructions had any influence on behavioral effects. Different from what was observed by Schmidt and De Houwer (2012a), explicit instruction did not enhance contingency awareness and had no impact on congruency effects in our paradigm.

Importantly, the reliable congruency effect in the color-only contingency blocks suggests the existence of a semantic component in color-word contingency learning. This was corroborated by the joint analysis, with more statistical power than the individual experiments.

## Experiment 3

In the preceding experiments, we used horizontally arranged color patches as response cues, such that each word’s print color could be matched directly with the colored response cues on the screen. In Experiment 3, we replaced the colored response cues by color names printed in black. Obviously, this will slow responding, because matching the print color of the stimulus word with one the four black words *rot, grün, gelb*, and *blau* necessitates at least one intermediate step: The retrieval of the German *name* of the pseudoword’s color. This name can be matched to one of the four color words present on the screen, whose positions correspond to the response buttons (see Figure 6). Thus, we expected prolonged response times, and predicted that the additional color-name retrieval step facilitates contingency learning.

**Figure 6.**
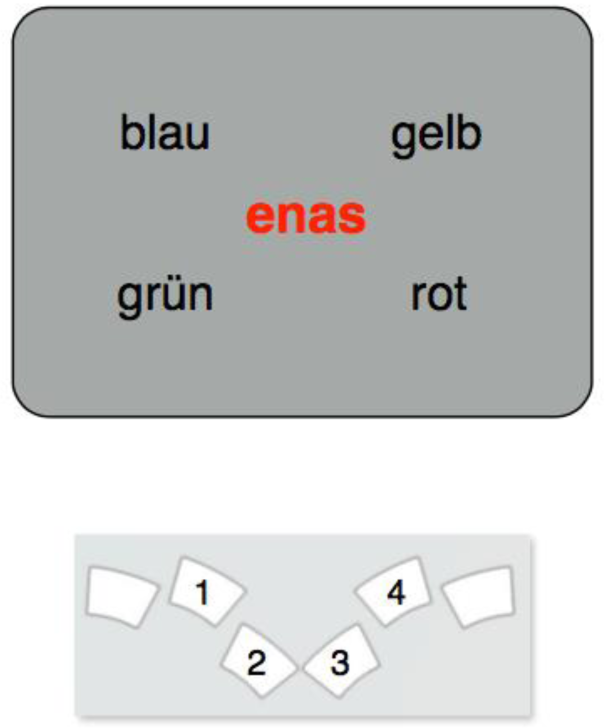
Stimulus display and spatial arrangement of response buttons in Experiment 3. As above, numbers on the response buttons are shown here to indicate which buttons were to be used. In the experiment, they were not actually labelled. In the sample trial shown here, response button 3 is the correct response.

### Methods

#### Participants

Nine students (6 male, age: *M* = 22.44, *SD* = 2.70) took part in Experiment 3. Inclusion criteria, informed consent and compensation were the same as above.

#### Materials and Procedure

Experiment 3 was a close replication of Experiment 1. The crucial change was that we replaced the color squares by their corresponding German color names (*rot, gelb, grün, blau*) in black print color. To minimize spatial position effects on reading times, these words as response cues were positioned in the corners of an invisible rectangle, centered on the fixation cross, thus equating their eccentricity (cf. Figure 6). To increase stimulus-response compatibility, the spatial arrangement of the response buttons approximated the rectangular display of the color words on the screen, allowing comfortable responding with the index und middle-fingers of the hands. As in Experiment 1, participants were not informed about the underlying contingencies. Except for the changes detailed above, all details of procedure and data analysis, including the matching task for assessing contingency awareness, were identical to those in Experiments 1.

### Results

Mean response times as well as net effects for Experiment 3 are shown in Figure 7. The pattern of response times again resembled those from Experiments 1 and 2. There was a clear and early-emerging congruency effect (of 28 ms) that was stable across contingency types and block repetitions. The ANOVA of the mean response times confirmed a significant main effect of *Congruency*, *F*(2, 16) = 24.64, *p* < .001. Responses were faster on congruent than on neutral trials (784 ms vs. 811 ms, *t*[8] = 5.69, *p* < .001), which did not differ from incongruent trials (812 ms, *t*[8] = 0.46, *p* = .654). Mean response times decreased significantly over *Block Repetitions*, *F*(7, 56) *=* 47.86, *p* < .001, from 956 ms in Block 1 to 729 ms in Block 8, and were affected by *Contingency Type*, *F*(3, 24) = 102.69, *p* < .001. Response speed did not reliably differ between *Col*- (940 ms) vs. *Pos*-blocks (938 ms), *t*(8) = 0.38, *p* = .744, but did so between *Pos*- and *ColPos*-blocks (696 ms), *t*(8) = 8.96, *p* < .001, as well as between the *ColPos*- and *ColPosFix*-blocks (633 ms), *t*(8) = 6.16, *p* < .001. None of the interactions turned out significant (all *F* < 1.00 and *p* > .441).

As in Experiment 1 and 2, we assessed the significance of individual congruency effects. The mean response time difference between congruent and incongruent trials was significant in all contingency conditions: *Col*: 29 ms (*t*[8] = 3.43, *p* = .009); *Pos*: 20 ms (*t*[8] = 2.42, *p* = .042), *ColPos*: 44 ms (*t*[8] = 3.72, *p* = .006) and *ColPosFix*: 19 ms (*t*[8] = 2.76, *p* = .025).

Errors rates were slightly higher than in Experiments 1 and 2, now 3.97% (congruent 3.60%, neutral 4.20%, incongruent 4.12%; see Supplementary Materials, Figure A1). The main effect of *Congruency* just failed significance, *F*(2, 16) = 3.60, *p* = .051. No other main effects or interactions reached significance, all *F* < 2.27, all *p* > .106.

**Figure 7.**
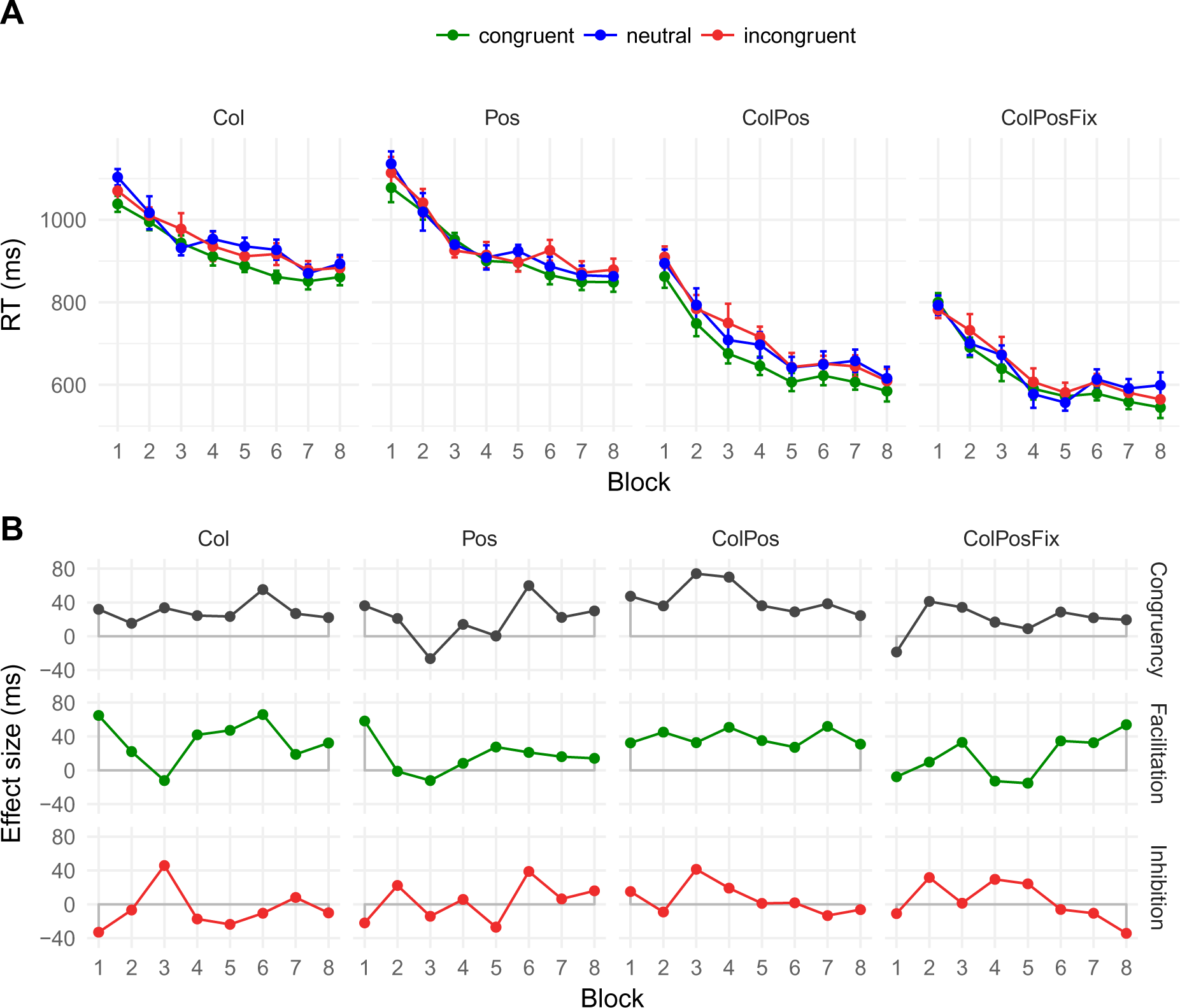
Mean response times (A) and net effects (B) in Experiment 3 (n = 9). Error bars and abbreviations as in Figure 3. Note the larger scale on the vertical axis, compared to the figures from Exp. 1 and 2.

#### Contingency Awareness

The color-word matching performance after session 4 differed substantially across contingency types (range: 16 - 47%, chance level = 25%). As in Experiment 1 and 2, words from *ColPosFix* blocks were matched to their corresponding colors with better than chance accuracy (M = 47%, SD = 20%, *t*(8) = 3.41, p = .005, BF_10_ = 13.47). In the remaining conditions, however, neither color- nor position-matching performance was better than chance (all *t*[8] < 1.49, *p* > .090) and either inconclusive or in favor of the null (BF_01_ between 0.74 and 5.86; see Supplementary Materials, Table A3, for details).

### Discussion

In Experiment 3, we tested whether replacing color *patches* by color *words* generates the same patterns of congruency effects. As expected, the response-time range clearly changed, now spread over a much larger interval (from 550 to 1100 ms), as compared to Experiments 1 and 2 (from 480 to 630 ms). Color-only and position-only contingencies in particular slowed responses. When both color and position correlated with stimulus words, however, correct responses were almost as fast as in Experiments 1 and 2.

Large increases in response time notwithstanding, the pattern of effects was similar to that in Experiments 1 and 2. Most importantly, there were robust congruency effects for all contingency types: from 19 and 44 ms in the two conditions with both color and position information to 20 ms for position-only and 29 ms for color-only information. The latter effect provides clear evidence for a semantic component in associative learning. Considering congruency effects within blocks, all but two out of the 32 conditions showed faster mean responses on congruent trials than on incongruent trials. This congruency effect was present early on, and there was no indication that it increased significantly across the eight blocks. That contingency-learning effects can arise so quickly, within a small number of trials, is evidence of the efficiency of associative learning. Experiment 4 provides additional, unintended support for its remarkable speed and plasticity.

## Experimenr 4

Logic and intention of Experiment 4 were the same as of Experiment 1. In fact, it was actually designed and run as the first experiment in this series. Only when we analyzed the data, did we discover a subtle programming bug which, however, turned out to provide valuable insights into the dynamics and stability of the implicit-learning processes.

The effect of the programming bug was that - different from what we had intended - the word-color and/or word-position contingencies did not remain constant throughout the experiment. Rather, although contingencies *within* any trial block were implemented as planned, they were re-randomized before each new block of trials. Thus, any word-color and/or word-response contingency that had been learned *within* a trial block could not facilitate responding in subsequent blocks – indeed, it should generate interference as in AB-AC experiments. Even though Experiment 4 was run first, we decided to present the originally intended Experiments 1 – 3 first, before revealing the findings from this deviating implementation.

### Methods

#### Participants

Thirteen students (4 male, age: *M* = 23.69, *SD* = 3.07) took part in Experiment 4. Inclusion criteria, informed consent and compensation were the same as above.

The experiment followed the same 4 x 3 x 8 experimental design, crossing *Contingency Type* (*Col, Pos, ColPos, ColPosFix*), with *Congruency* (congruent, incongruent, neutral) and *Block Repetition* (1 to 8). Within each block of 75 trials, a fixed subset of five pseudowords was used: one neutral control word and four experimental words of the same type (either *Col, Pos, ColPos, or ColPosFix*). Each word appeared as a colored stimulus on 15 trials per block.

Procedural details were identical to those of Experiment 1, except for the following change: For each participant, five pseudowords were randomly assigned to each of the four contingency types. However, this assignment *applied to the current trial block only*; before the next trial block of that condition the five words were *randomly reassigned* to color and/or position. Thus, the mapping of pseudowords to particular colors or response-positions was re-shuffled from one block to the next. Unknown to the participants (and to the experimenters), any implicit associations from pseudowords to colors, response positions, or their conjunction that had been established within any one block became useless or even detrimental in the next trial block of that condition.

Remarkably, none of the participants reported having noticed any change of stimulus-response contingencies between trial blocks and sessions. In fact, we discovered the programming bug only when we (disappointed by the absence of any learning effects in the overall data) scrutinized the stimulus generating programs. However, because stimulus-response contingencies were as planned *within*, but randomly re-assigned *between blocks*, we checked the data for traces of implicit learning *within* trial blocks. To do so, we subdivided each block into five non-overlapping segments (‘sub-blocks’) of 15 trials each and checked whether and how mean RTs and error rates (averaged over all eight repetitions of a given contingency condition) changed *within* blocks. Moreover, the final segments of the blocks were analyzed in a repeated-measures ANOVA with factors *Congruency* and *Contingency Type*.

### Results

In all contingency conditions, responding in the last segment of blocks was faster in the congruent than in the incongruent condition, by 10 ms on average (see Figure 8). Analysis of mean response times in this last segment revealed main effects of *Congruency* (*F*(2, 24) = 5.46, *p* = .011), and of *Contingency Type* (*F*(3, 36) = 7.43, *p* < .001), but no reliable interaction (*F*(6, 72) = 0.92, *p* = .486). For the mean overall 10 ms congruency effect, pairwise *t*-tests showed that responses on congruent trials (543 ms) were slightly but not significantly faster than those on neutral trials (547 ms), *t*(12) = 1.61, *p* = .132, whereas responses were faster on neutral than on incongruent trials (553 ms), *t*(12) = 2.38, *p* = .035.

Error rates in the final block segment were between 1.7% and 3.6% but showed the expected pattern only in the Pos and ColPosFix conditions. None of the main and interaction effects turned out reliable in the repeated-measures ANOVA on the arcsine square-root transformed error rates (*Contingency Type: F*(3, 36) = 0.24, *p* =.868; *Congruency: F*(2, 24) = 1.47, *p* = .250; *Contingency Type* × *Congruency*: *F*(6, 72) = 0.98, *p* = .445).

**Figure 8.**
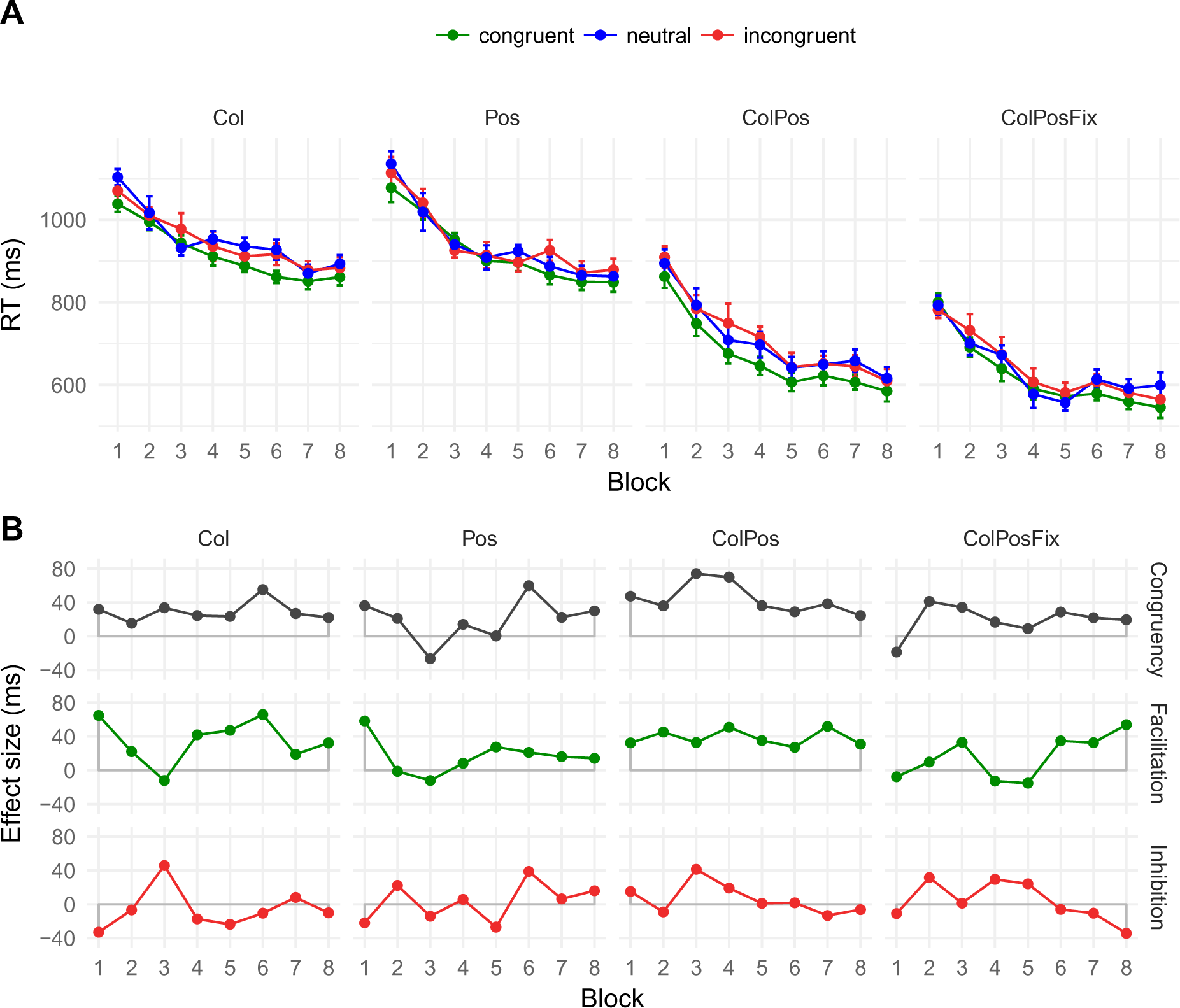
Mean response times (A) and net effects (B) in Experiment 4. Error bars and abbreviations as in Figure 3.

### Discussion

Experiment 4 assessed whether there are traces of implicit learning even when the contingencies between stimulus words and their print color and/or correct response position are randomly reshuffled from block to block. Indeed, our results show that this is the case: Responses in the final block segments yielded an overall 10 ms congruency effect. There was no evidence that the size of the congruency effect was modulated by contingency type, but a differentiation is difficult given the limited number of learning trials for any word-color or word-position pairing. Nevertheless, the data from Experiment 4 reveal that, in spite of the task complexity and the non-unintended changes of the stimulus-response contingencies from block to block, color-word contingency learning can swiftly lead to congruency differences in response times, even within single blocks that contain 12 congruent and 3 incongruent trials only for any given pseudoword.

#### General Discussion

In four experiments, we investigated the mechanisms underlying color-word contingency learning in the color-matching Stroop task. Our goal was to disentangle potential semantic and response-learning contributions to the learning effect. We employed a novel variant of the paradigm in which a row of color patches (Experiments 1, 2, and 4) or a set of color words printed in black (Experiment 3) on the screen indicated which of four response buttons corresponded to which of the four print colors on a given trial. This task design allowed us to independently manipulate word-to-color and word-to-response-position contingencies. In fact, we implemented separate contingency conditions in which different pseudowords were associated either with print color (*Col*), with spatial position of the assigned response (*Pos*), or with both (*ColPos*). In the fourth contingency type (*ColPosFix*), the pseudowords were also associated with both color and response position, while keeping the spatial order of the response color patches/words constant throughout the experiment. This latter condition most closely resembles the situation in previous experiments with a fixed set of colored response buttons. Importantly, participants adapted to the particular contingency – and thus learned - with all contingency types.

Overall, the response-time gains and losses caused by the different contingency types were small but statistically reliable. With color patches as response cues (Experiments 1 and 2), the strongest congruency effects were seen in conditions that contained position contingencies (effects between 8 and 13 ms), but the color-contingency-only condition also revealed a small semantic (word-color) learning effect (5 ms). With color words printed in black as response cues (Experiment 3), effects were substantially larger and more similar across the four contingency types (around 30 ms on average). Remarkably, the evidence for inhibition on incongruent trials (relative to neutral ones) was weak or absent: The net inhibition RT effects across contingency types were a mere 3, 3, and 1 milliseconds in Exps. 1, 2, and 3, respectively, which stands in marked contrast to the corresponding net facilitation effects of 11, 8, and 27 ms. This difference in effect size is likely due both to the rare occurrence of incongruent trials (20% of trials) and the fact that, within each trial block, the incongruent trials involved each correct response position and/or color just once.

Our experimental design was much more complex than in previous color-word contingency learning studies (but see Schmidt et al., 2018). The different types of contingency were manipulated within-participants, in separate blocks, with a neutral control word in each contingency condition. Throughout the experiment, participants were to process 20 novel words, and for 16 of these, contingencies existed that could be learned. We also randomly varied the font type and size of the pseudowords from trial to trial, to exclude mere context-specific, visual learning effects (cf. Schmidt & Lemercier, 2018). Finally, when response cues alternated positions on a trial-by-trial basis, participants had to locate the correct response-button position first before executing the color-matching response. This certainly increased task demands, as compared to responding with color-labelled physical buttons. In spite of these design and stimulus characteristics, we obtained reliable and consistent congruency effects across contingency conditions and experiments.

The observation (in Experiments 1 and 2) that congruency effects were larger for response-contingency learning than for semantic learning replicates findings from the 2:1 mapping paradigm used to distinguish semantic and response learning (Schmidt et al., 2007). Schmidt and colleagues reported a sizeable 26-millisecond congruency effect as evidence of response learning. Because there were no additional stimulus-match effects, Schmidt and colleagues concluded that semantic learning contributes little to color-word contingency learning. This is different in our study, however: Participants did profit from semantic associations that existed in the color-contingency-only blocks. Our results clearly indicate that even such an incidental statistical association of words and colors is sufficient to foster semantic learning. Our use of semantically neutral pseudowords rather than of meaningful words may well have facilitated semantic learning, as it is probably easier to learn semantic features for novel words such as “enas” that have no semantic representation, than for existing words such as “month” that have a semantic representation but – for most of us – are non-specific for color. Finally, with color words rather than color patches as response cues (Experiment 3), semantic effects were as large as pure response-position effects, and as the effects of color and position combined.

Interestingly, our results provide no evidence for additivity of position and color contingencies. Purely semantic (*Col* conditions) and response-learning effects (*Pos* condition) do not add up to substantially larger effects in the combined conditions (*ColPos* and *ColPosFix*). This was most obvious in Experiment 3, where the pure semantic and response-learning effects were relatively large and about equal in size (ca. 30 ms); however, the effects in the combined blocks did not increase, as would be expected from additive accounts.

Next, in all four experiments, contingency effects emerged early on and did not reliably increase with repetition. The inadvertent data from Experiment 4 corroborate this: even when word contingencies are randomly reset from block to block, 12 congruent trials for any word seem to suffice to yield contingency adaptation. These findings are fully consistent with those of Schmidt et al. (2010), a single-session study. However, our findings contrast with studies in which *shapes* rather than *words* were associated with colors. Under these conditions, interference effects in color naming emerged after intensive training only (MacLeod & Dunbar, 1988).

An important insight from our experiments concerns contingency awareness. Experiments 1 and 2, with color patches as response cues, differed with respect to instructions only; in Experiment 2, but not in Experiment 1, participants were informed about the global type of contingency present in each upcoming block (color, position, or both). This explicit information had no impact on response times, however, nor on the contingency-awareness data collected afterwards. We were thus unable to induce contingency awareness to the extent that it showed up in the post-learning assessment. This contrasts with results of Schmidt and De Houwer (2012a), who observed significantly larger congruency effects when participants were informed about, and indeed aware of, the type of contingency, as when not. Note that their experiment included only one type of contingency and only three word stimuli, which certainly may have facilitated focusing on this one type of association, as compared to our more complex experiment.

In sum, our findings suggest two important properties of the underlying learning process:

1. This type of learning is powerful: In spite of the incidental and actually task-irrelevant associations with which words are presented, participants need just a few trials to optimize behavior to the additional information provided by the color and/or response contingencies.
2. The impact of these learned contingencies on response time does not depend on contingency awareness, and seems limited: congruency effects do not substantially increase with block repetitions. This is particularly surprising given the swift initial learning. It may be that the incongruent and neutral trials within each block, along with the conflicting information from other blocks (e.g., other words associated with the same colors/positions), discourage participants from relying more strongly on the learned associations.

How do our results relate to previous studies on semantic novel-word learning and to language learning in general? Most previous studies that investigated semantic learning from statistical association used richer semantic input and employed more explicit learning paradigms (e.g., Breitenstein et al., 2007; Dobel et al., 2010; Geukes et al., 2015; Yu & Smith, 2007). Here, participants merely had to match the pseudoword’s ink color to the corresponding color patch or word, which required no attention to the pseudoword itself. Indeed, the contingency-awareness task showed that participants matched words from the *Col* blocks no better than chance, suggesting that no explicit word-color links were available. Interestingly, the only evidence for above-chance matching was found in the *ColPosFix* condition that mimicked the classic color-word contingency learning experiment with confounded color and position learning (Schmidt et al., 2007). Based on the participants’ report of noticing the contingency or not, Schmidt and colleagues distinguished between subjectively aware and unaware participants, but they did not formally test any matching performance. Thus, despite of not being able to report the contingencies, their participants might have been able to match words to colors with above chance accuracy.

Although the task in our experiments did not require any semantic processing and there was no explicit role for semantics (in contrast to previous more explicit word-learning experiments) we observed a semantic effect (similar in nature to the effect from our Stroop study with pseudowords, Geukes et al., 2015). Thus, even associations presented in a purely incidental fashion, largely out of our attention’s focus, seem to be monitored constantly to improve our repertoire of word-meanings links.

Given that sizeable semantic effects of even 30 ms were obtained in a highly complex design with four different contingency types, it seems worthwhile to replicate this finding in a simpler experimental context, focusing on the *Col*-blocks. It might also be worthwhile to test whether novel “color words” learned during color-word contingency learning are sufficiently associated with their color meanings as to produce more indirect semantic effects, such as behavioral semantic priming, or an N400 reduction in event-related potentials. Such effects would be directly comparable to more general semantic learning effects observed by Breitenstein et al. (2007), Dobel et al. (2010), and Geukes and Zwitserlood (2016).

In summary, our variant of color-word-contingency learning with color cues on screen is a suitable tool for separating semantic and response-learning components. Our findings confirm previous studies that reported clear evidence for response learning in color-word-contingency learning. Over and above those findings, we here demonstrated unequivocal semantic learning effects, showing that incidental learning via statistical association can be sufficient to establish effective word-meaning links.

## Acknowledgements

We are grateful to Michael Praast, Britta Radenz, Lukas Urban, and Dario Zaremba for their help in conducting and analyzing the experiments in this study. We also thank Jens Bölte for helpful discussions.

## Footnotes

i *Stroop congruency effect* here refers to the finding that indicating the print color of a word (via button press) is faster and less error-prone when the word itself refers to its print color (e.g., *red* printed in red) rather than to a different color (e.g., red printed in blue). Under the conditions stated above, we observed similar, albeit smaller Stroop effects (compared to the native-language Stroop effect) for novel pseudowords that had been associated with the native language color words (e.g., *binu* associated with red; Geukes, Gaskell & Zwitserlood, 2015).

ii Note that our task differs from the standard Stroop matching task, which is essentially a Yes-No (match/mismatch) procedure. In contrast, the four-color matching task, first used by Schulz & Liebing (1991), avoids the methodological and theoretical pitfalls of the match/mismatch procedure pointed out by Luo (1999) and Goldfarb & Henik (2006). More importantly, the four-color matching task allowed us to independently manipulate color-word and color-response contingencies, as will be detailed in the following.

iii The Bayes-Factor of BF_10_ = 3.76 indicates that the observed pattern is almost four times as likely under the alternative hypothesis (‘performance better than chance’) than under the null hypothesis (‘performance at chance level or worse’). “Conventional approximate guidelines for strength of evidence were provided by Jeffreys (1939), though Bayes factors stand on their own as continuous measures of degrees of evidence. If BF > 3 then there is substantial evidence for H1 rather than H0; if BF < 1/3 then there is substantial evidence for H0 rather than H1; and if BF is in between 1/3 and 3 then the evidence is insensitive” (Beard, Dienes, Muirhead, & West, 2016).

