## Supplement Figure&Tables for "Disentangling semantic and response learning effects in color-word contingency learning"

*Supplementary Materials*

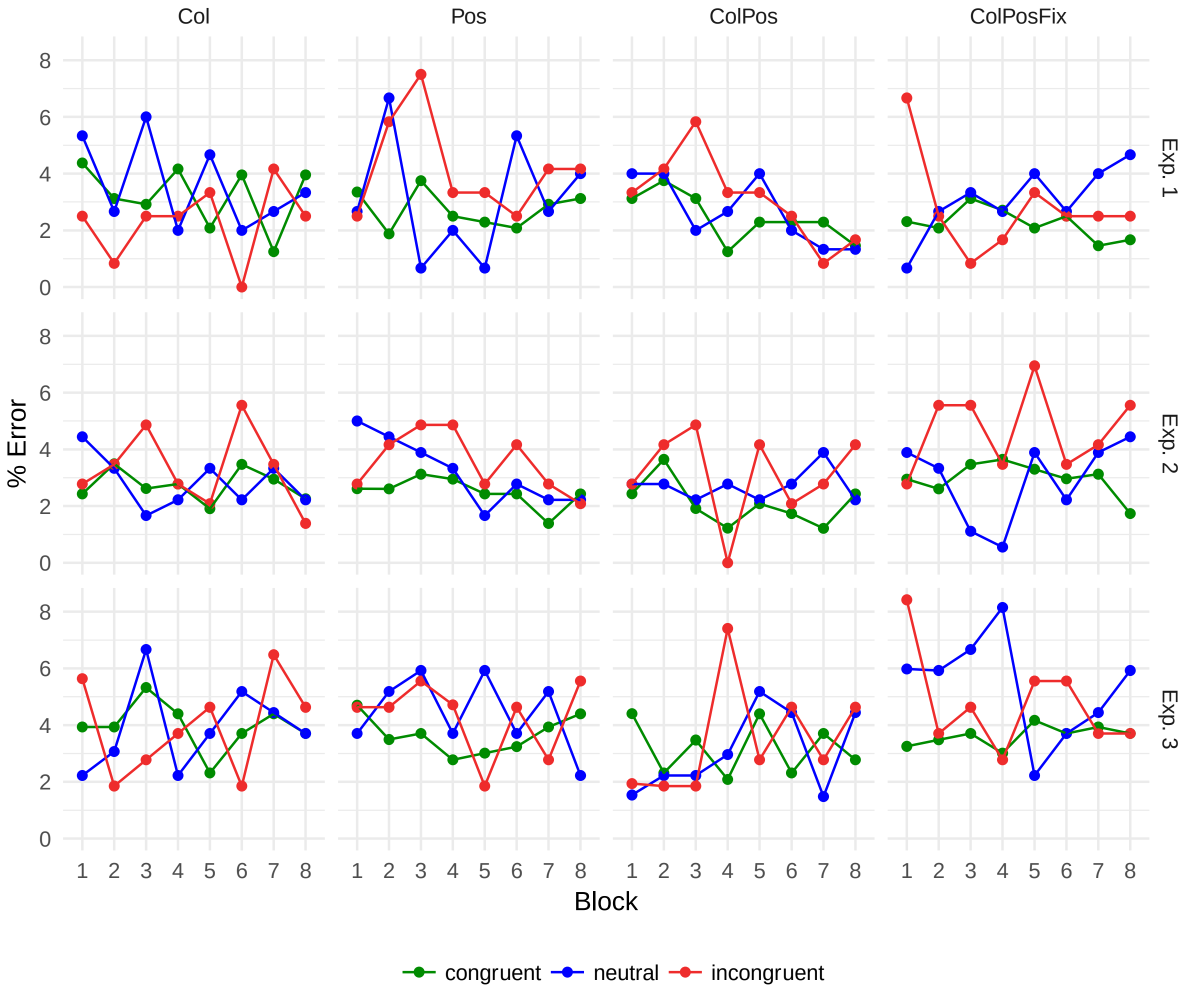


**Figure A1.** Mean error rates in Experiments 1, 2, and 3.

*Table A1. Matching performance and Inferential statistics of in Experiment 1.*

| word type | *t(9)* | *p* | $BF_{10}$ | $BF_{01}$ |
|---|---|---|---|---|
| Matching by color | | | | |
| Col | 1.63 | .068 | | 1.19 |
| ColPos | 1.41 | .096 | | 1.49 |
| ColPosF | 2.86 | .009 | 3.76 | |
| Matching by position | | | | |
| Pos | 0.43 | .338 | | 2.99 |

| | | | |
|---|---|---|---|
| ColPos | 1.91 | .045 | 1.15 |
| ColPosF | 3.10 | .007 | 5.11 |

*Note: t-tests are one-sample t-tests testing whether participants' matching of the respective four words to colors or positions was better than chance level (25%): Accordingly, one-sided p-values are reported. Scaled JZS Bayes Factors are also given, based on an r scale parameter of 0.707. $BF_{10}$ denotes the Bayes Factor in favor of the alternative hypothesis (if $BF_{10} > 1$, evidence is in favor of above chance performance). When $BF_{10}$ is below 1, that is, when evidence is in favor of the null hypothesis (chance performance or worse), the reciprocal $BF_{01}$ (= 1/ $BF_{10}$) is given instead, to facilitate interpretation.*

### *Table A2. Matching performance and inferential statistics in Experiment 2.*

| word type | *M (SD)* | *t(11)* | *p* | $BF_{10}$ | $BF_{01}$ |
|---|---|---|---|---|---|
| Matching by color | | | | | |
| Col | .23 (.14) | 0.80 | .219 | | 1.73 |
| ColPos | .25 (.17) | 1.00 | .169 | | 1.41 |
| ColPosF | .42 (.26) | 2.86 | .008 | 8.23 | |
| Matching by position | | | | | |
| Pos | .28 (.23) | 1.24 | .121 | | 1.08 |
| ColPos | .18 (.18) | -0.32 | .623 | | 4.32 |
| ColPosF | .20 (.21) | 0 | .500 | | 3.48 |

*Note that neutral words were included in the matching task in Exp. 2 and chance level is thus at 20%.*

### *Table A3. Matching performance and inferential statistics in Experiment 3.*

| word type | *M (SD)* | *t(11)* | *p* | $BF_{10}$ | $BF_{01}$ |
|---|---|---|---|---|---|
| Matching by color | | | | | |
| Col | .31 (.30) | 0.56 | .297 | | 1.98 |
| ColPos | .33 (.35) | 0.71 | .250 | | 1.72 |
| ColPosF | .47 (.20) | 3.41 | .005 | 13.47 | |
| Matching by position | | | | | |
| Pos | .25 (.19) | 0 | .500 | | 2.97 |
| ColPos | .41 (.30) | 1.49 | .090 | 1.35 | |
| ColPosF | .16 (.19) | -1.43 | .901 | | 5.86 |

*Note. As in Exp. 1A, chance level is at 25%.*

### *Table A4. Summary statistics for all four experiments.*

| | Col | | | Pos | | | ColPos | | | ColPosFix | | |
|---|---|---|---|---|---|---|---|---|---|---|---|---|
| exp. | mean | s.d. | CE | mean | s.d. | CE | mean | s.d. | CE | mean | s.d. | CE |
| 1 | 586 | 21 | 4 | 593 | 24 | 14 | 547 | 25 | 15 | 530 | 24 | 15 |
| 2 | 618 | 22 | 7 | 618 | 25 | 8 | 576 | 28 | 17 | 546 | 33 | 11 |
| 3 | 940 | 76 | 29 | 938 | 71 | 20 | 696 | 68 | 44 | 633 | 61 | 19 |
| 4 | 561 | 17 | 8 | 562 | 20 | 5 | 528 | 22 | 12 | 529 | 24 | 15 |

*CE = net congruency effect RTincongruent - RTcongruent (Exp. 1-3: across all blocks; Exp. 4: last block segment, averaged across all blocks)*
